# High Frequency MHz-Order Nanovibration Enables Cell Membrane Remodelling and Lipid Microdomain Manipulation

**DOI:** 10.1101/2024.09.24.614713

**Authors:** Lizebona A. Ambattu, Blanca del Rosal Rabes, Charlotte E. Conn, Leslie Y. Yeo

**Affiliations:** Micro/Nanophysics Research Laboratory, School of Engineering, RMIT University, Melbourne, VIC 3001, Australia; School of Science, RMIT University, Melbourne, VIC 3001, Australia

## Abstract

We elucidate the mechanism underpinning a recently discovered phenomenon in which cells, quite unexpectedly, respond to MHz-order mechanostimuli. Deformations induced along the plasma membrane under these external mechanical cues are observed to decrease the membrane tension, which, in turn, drives transient and reversible remodelling of its lipid structure. In particular, the increase and hence coalescence of ordered lipid microdomains leads to closer proximity to mechanosensitive ion channels—Piezo1, in particular—that due to crowding, results in their activation to mobilise influx of calcium (Ca^2+^) ions into the cell. It is such modulation of this second messenger that is responsible for the downstream signalling and cell fates that ensue. Additionally, we show that such spatiotemporal control over the membrane microdomains in cells—without necessitating biochemical factors—facilitates aggregation and association of intrinsically disordered tau proteins in neuroblastoma cells, and their transformation to pathological conditions implicated in neurodegenerative diseases, thereby paving the way for the development of therapeutic intervention strategies.

## Introduction

The plasma membrane—formed from the self-assembly of amphiphilic lipids, transmembrane and membrane-anchored protein clusters, and carbohydrates—comprises the physical boundary that separates and regulates the interactions between the interior of a cell from its external environment. Due to lipid–lipid and lipid–protein interactions, liquid–liquid phase separation results in the heterogeneity of these lipid and protein assemblies, which manifest as distinct microscopic lateral clusters (>300 nm). These clusters can be separated into relatively ordered liquid domains (L*_o_*) known as lipid rafts that contain saturated lipids (phospholipids, glycolipids, and predominantly, sphingolipids) and sterols (mainly cholesterol) (*1–4*), and disordered clusters (L*_d_*) containing unsaturated lipids, with transmembrane proteins partitioned between both regions (*5*). It is the dynamic spatial reorganisation of these regions in response to alterations in the membrane tension as a consequence of deformations imposed on the plasma membrane that enable its role as a primary mechanosensor (*6–8*), which, in turn, plays a crucial role in signal transduction activation that is the determinant of downstream cellular fate.

Despite their biological significance, the ability to directly manipulate the lipid microdomains on the plasma membrane—which could not only provide the elusive tool to study lipid rafts but also enable a potent means for engineering cells—remains challenging without modifying lipid composition, osmolarity, or the underlying substrate, or inducing vesicle fusion, all of which fall short of representing key biological processes that recapitulate typical *in vivo* conditions (*9*). While local heating with a laser allows the possibility for inducing reversible microdomain transition between the L*_o_* and L*_d_* phases, this has only been demonstrated in *model lipid bilayers* (i.e., vesicles) (*10*).

Recently, we had observed that cells exhibit specific responses when subject to high frequency (*O*(10 MHz)) mechanostimulation (*11*), which, in itself, is surprising* given that such frequencies are far beyond the typical perception range (several Hz, commensurate with that associated with physiological motion) expected of cells. Central to all of these responses and the downstream cell fates they induce (e.g., cell adhesion, migration and proliferation (*13, 14*), vesicle trafficking and exosome biogenesis (*15*), immune cell activation (*16*), stem cell differentiation (*17, 18*), and endothelial barrier modulation (*19*)) are the transcriptomic changes brought about by second messenger signalling that is instigated by the ability for the external high frequency mechanical cues to modulate Ca^2+^ levels within the cell through the activation of various mechanosensory elements along the plasma membrane, such as mechanosensitive ion channels and membrane-associated proteins (*11*).

In this work, we report our attempt to uncover the fundamental mechanisms responsible for this signal transduction process. In doing so, we shed light on the processes by which the high frequency mechanostimulation, applied for just several minutes, drives transient deformation of the cell and reorganisation of its cytoskeletal structure through a dynamic combination of compression, tension and shear. In particular, we show that the resultant change in the cell’s membrane tension as a consequence leads to spatiotemporal remodelling of the lipid structure and hence changes in its microdomain distribution.

Parenthetically, we note that the phenomena we report here due to 10-MHz-order mechanostimulation, in the form of nanometre amplitude surface acoustic waves (SAWs; typically >20 MHz) and, hybrid surface and bulk acoustic waves, i.e., surface reflected bulk waves (SRBWs; typically <20 MHz) (*20*), are considerably distinct from that reported with 1 kHz order vibration (*21–23*), and, more generally, any mechanostimuli conducted at frequencies below 1 MHz using focussed ultrasound (*24*). Given that cavitation, which is notably absent at 10 MHz order frequencies under the intensities we typically employ (*25*), becomes increasingly prevalent at lower frequencies, it is likely that prolonged exposure leads to disruption of the lipid microdomains due to pore formation along the membrane (*24, 26*). As such, the effects we observe in the present work due to the high frequency mechanostimulation, particularly its ability to directly manipulate the lipid microdomains on the plasma membrane, do not appear to have been reported at lower frequencies. Additionally, we note that the observed phenomena in this work cannot be attributed to flow-induced shear since the intensities we typically employ are below that required to generate appreciable acoustic streaming (i.e., the flow arising as a consequence of viscous dissipation as the sound waves attenuate in the liquid medium).

Motivated by our observations that the spatiotemporal lipid microdomain manipulation arising as a consequence of the high frequency mechanostimulation facilitates activation of ordered membrane-associated proteins such as Piezo1, we also show in the final section of this work, as an example to highlight its potential, the intriguing possibility for transforming intrinsically disordered proteins—in particular tau (tubulin-associated units), whose function and structure depend on their interactive partners, membranes, proteins or RNA, and whose association with the plasma membrane and aggregation has been implicated in cell senescence in the brain, and hence the pathology of neurodegeneration (e.g., Alzheimer’s disease and and frontotemporal dementia (*27–29*)). This demonstration of mechanical manipulation of intrinsically disordered proteins through membrane microdomain reorganisation provides a means for understanding the complex signalling processes accompanying the association of these proteins to the plasma membrane during pathological transformation that occurs during the progression of neurodegenerative diseases, therefore paving the way for the development of interventional strategies to treat these diseases.

## Results and Discussion

As model cell lines, human mesenchymal stem cells (hMSCs) from bone marrow and SHSY5Y neuroblastoma cells cultured in glass-bottom cell culture plates were exposed to 1.5 and 2.5 W SRBW, as shown in the experimental setup illustrated in Fig. 1(a); these powers were chosen based on optimisation studies that reveal the most significant mechanoresponse for these particular cells. We then probe changes in their membrane tension and lipid structure at different post-incubation periods using various assays, as described in the Methods section.

**Figure 1.**
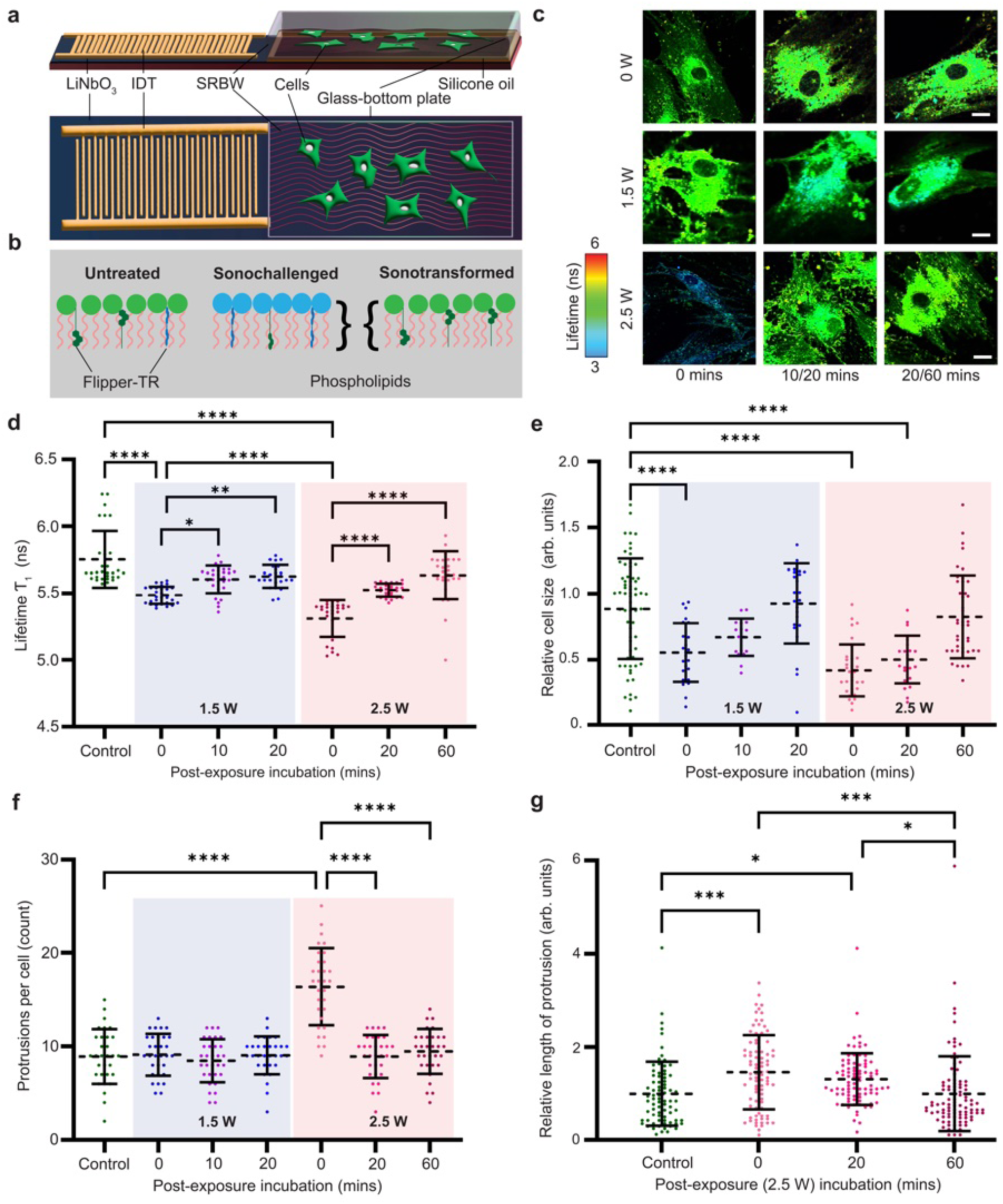
*(preceding page)*: **Experimental setup and plasma membrane tension changes in response to the SRBW forcing**. **a** Top and side view schematics showing the experimental setup in which the SRBW (not to scale), generated along a piezoelectric lithium niobate (LiNbO_3_) substrate by applying an AC electric signal at the device’s resonant frequency (10 MHz) to an interdigitated transducer electrode (IDT) photolithographically patterned on the substrate, is coupled through a thin layer of silicone oil into a glass-bottom cell culture plate containing the adherent cells. **b** Schematic representation of the changes in the plasma membrane tension due to the SRBW irradiation, measured using the probe Flipper-TR™, which when tagged to the membrane lipids, allows quantification of the pressure along its axis: ‘sonochallenged’ cells (i.e., the state of the cells immediately following cessation of the SRBW irradiation (0 mins post-exposure incubation)) exhibited a reduced fluorescence lifetime similar to cells under hyperosmotic conditions, while ‘sonotransformed’ cells (i.e., the state of the cells following their recovery (20–60 mins post-exposure incubation)) regained higher fluorescence intensity lifetime values close to that of untreated (control) cells owing to relaxation of the membrane. **c** Representative images (colour-coded for the overall fluorescence lifetime) of Flipper-TR™probed hMSCs exposed to 8 mins of SRBW irradiation at input powers of 0, 1.5 and 2.5 W and fixed at different post-exposure incubation durations; control untreated cells (0 W) were fixed at 0, 20 and 60 mins, while cells irradiated at 1.5 and 2.5 W were fixed at 0, 10 and 20 mins, and 0, 20 and 60 mins, respectively, determined through a series of optimisation studies. Scale bars denote a length of 40 µm. Comparison of **d** the longest component of the fluorescence lifetime (intensity-weighted) ⌧_1_, **e** the cell area, **f** the average number of protrusions per cell, and, **g** the relative filipodia length in untreated and SRBW-irradiated cells; the data are represented in terms of its mean *±* the standard error over triplicate runs, and the asterisks *^⇤⇤⇤^* and *^⇤⇤⇤⇤^* indicate statistically significant differences with p < 0.001 and p < 0.0001, respectively.

### Deformation driven changes to the plasma membrane tension

Given the considerably longer SRBW wavelength at 10 MHz (A = 398 µm) but much smaller amplitude *O*(10 nm) compared to typical cellular dimensions, and as the SRBW vibrational forcing time scale (1/f *⇠ O*(10*^-^*^7^ s)) far exceeds characteristic cellular time scales (*O*(1–100 s) (*30*)), it would be reasonable to expect that cells are unlikely to deform under such high frequency mechanostimuli, at least under leading order modes and over time scales commensurate with the inverse of the frequency. Yet, considerable deformation is observed, as evident from the changes to the cells’ membrane tension observed, when they are exposed to the SRBW. Figure 1c,d, for instance, shows a marked decrease in membrane tension immediately upon exposure to the SRBW at input powers of 1.5 and 2.5 W, compared to that for unexposed (control) cells, as captured through the fluorescence lifetime of the membrane tension probe Flipper-TR™—a planarisable push–pull probe which, once inserted into the membrane lipid, undergoes conformable (planar vs twist) changes that influence the lifetime of its excited state (Fig. 1b)—bound to the cells. Specifically, the tension in the plasma membrane correlates linearly with the longest component of the fluorescence lifetime ⌧_1_, which, in turn, is dependent on the intramolecular charge transfer that occurs during changes in the twist angle and polarisation between the two twisted dithienothiophenes of the mechanophore (*31–33*). Owing to the short incubation periods in our experiments (<2 hrs), the ⌧_1_ values measured correspond solely to effects arising from the plasma membrane tension and not those due to changes in the intracellular structures. Importantly, we observe the mechanostimulated cells to eventually relax back to their original state; the larger the applied SRBW power, the longer the period over which this occurs, with the cells exposed to 1.5 W recovering within 20 mins and those exposed to 2.5 W recovering within 60 mins. This transient effect is consistent with observations in our previous studies, be it for intracellular delivery wherein such transient effects provided strong evidence, among others, that the SRBW did not result in physical pore formation, unlike in sonoporation (*34, 35*); or with endothelial cells, where barrier function was observed to be uniquely recovered following its initial permeabilisation following the sonochallenge—such recovery not being observed in, for example, with chemical insults to cells (*19*).

The decrease in membrane tension as a consequence of the SRBW mechanostimulation is akin to that observed in cells under hyperosmotic conditions (*32, 36, 37*); we note that the levels to which the membrane tension decreases due to the SRBW forcing, as captured by the reduction in ⌧_1_ by approximately 0.5–1 ns, is similar to that reported for these cells (*32*), or those mechanostimulated by static compression (*32,38*). In both these cases (i.e., cells exposed to hyperosmotic conditions or static compression), distinct morphological traits, such as contraction of the cell, a two-dimensional drum-like appearance on the apical side of the cell, and protrusions that resemble filopodial structures, were reported, all of which have been observed with cells exposed to the SRBW mechanostimulation (*34*). Given that these morphological traits appear to form immediately following the cessation of the stimulation, it is likely that they arise due to the retraction of the membrane towards the centre of the cell or nucleus, as indicated by the transient yet quick reduction in cell size (Fig. 1e), the increase in the number and length of protrusions (Fig. 1f,g), and the distribution of the membrane stain distinctly around the nucleus as a response to its contraction under the mechanostimulation, which can be seen to decrease towards the cell periphery (Fig. 1c). In any case, all of the characteristics observed can be seen to be more prominent at the higher SRBW intensity (i.e., 2.5 W), but in a manner akin to the recovery of the membrane tension to levels associated with its original state at rest, diminishes with post-exposure incubation time (Fig. 1c).

### Effect of changes in membrane tension on membrane fluidity

In contrast to cellular time scales, characteristic membrane lipid relaxation times (*O*(1 ps) (*39*)) are much shorter than the time scale associated with the SRBW forcing. As such, and given that the deformation-induced decrease in membrane tension is likely to alter the membrane fluidity and hence the homogeneity of its domain composition, it is not unreasonable to then expect modifications to the lateral organisation of the membrane lipid structure to accompany the SRBW mechanostimulation process. Indeed, we observe in Fig. 2 that the SRBW mechanos-timulated cells, stained with C-Laurdan—a commonly-used probe for determining membrane lipid order, display a notable increase in their generalised polarisation (GP) values^†^, which is indicative of an increase in the relative proportion between the ordered L*_o_* and disordered L*_d_* phases (*40*) (the L*_o_*/L*_d_* ratio being characteristic of the fluidity of the membrane and can vary between cell types and passages). More specifically, we note from Fig. 2b a shift toward higher GP values as the plasma membrane transitions towards a more homogeneous gel-like state with increasing SRBW power, compared to the unexposed control which exhibited a more heterogeneous composition comprising both L*_o_* and L*_d_* domains. Consistent with our earlier observations for the membrane tension though, we observe in Fig. 2c,e the peak distribution to shift back to values corresponding with the cells’ ground state with time: the heterogeneity in the domain composition with both low and high GP values can be seen to reappear within 60 mins post-incubation exposure for the SRBW mechanostimulated cells at 2.5 W.

**Figure 2.**
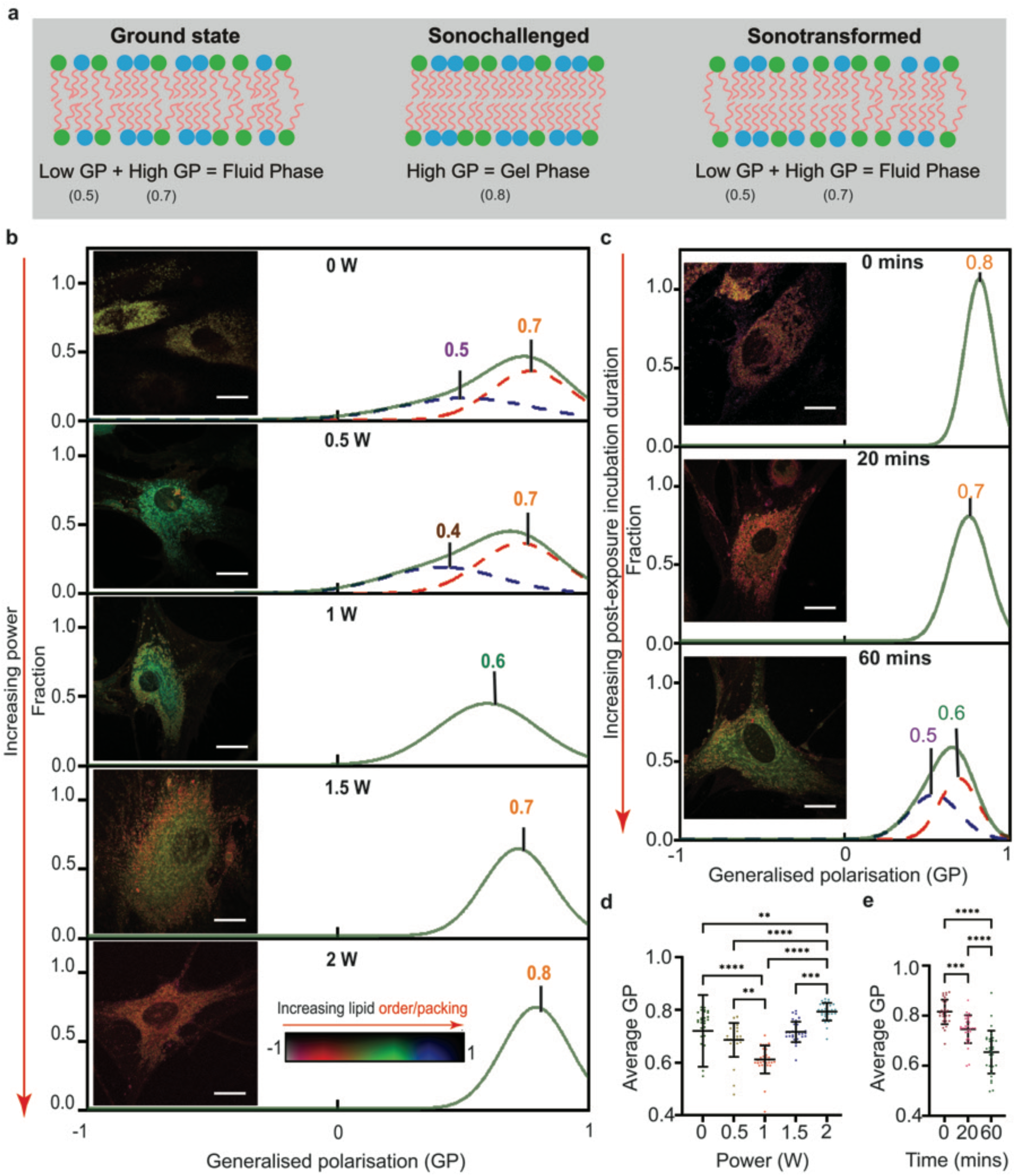
C**h**ange **in membrane fluidity due to the SRBW irradiation**. Schematic representation showing the effect of the SRBW on membrane fluidity: cells at ground state (and untreated cells) in which both L*_o_* and L*_d_* domains coexist (left), giving rise to both low and high generalised polarisation (GP) components; increased membrane fluidity in sonochallenged cells result in a transition to a more gel-like phase and hence higher GP values (centre); at longer post-exposure durations, sonotransformed cells return towards their ground state through increased exocytosis (right). **b**,c Images (inset) and distribution of their GP values (*-*1 to +1; as represented by the colour bar) for control and SRBW-irradiated C-Laurdan-labelled hMSCs, **b** fixed at the end of the exposure period, as a function of the SRBW power, and, **c** fixed at different post-exposure incubation periods for a SRBW power of 2.5 W. The histograms for the GP values were fitted with Gaussian distributions, with the solid green curves representing the average between low (dashed blue curves) and high (dashed red curves) distributions. Scale bars denote a length of 30 µm. **d**,**e** Average GP values in the cases shown in **b** and **c**, respectively.

The data are represented in terms of its mean *±* the standard error over triplicate runs, and the asterisks *^⇤⇤^*, *^⇤⇤⇤^* and *^⇤⇤⇤⇤^* indicate statistically significant differences with p < 0.01, p < 0.001 and p < 0.0001, respectively.

The increase in GP values, and hence membrane order and packing, with the SRBW mechanostimulation is, however, contrary to that observed for cells subject to hyperosmotic treatment, where decreases in membrane tension led to a decrease in L*_o_* domains (*32*). Similarly, our observations here are also contrary to that reported with increasing temperatures (*40*), therefore suggesting that the effect is unlikely a consequence of heating (which, in any case, is minimal given the low SRBW powers and short exposure duration of just a few minutes), consistent with our prior studies that verify that the SRBW mechanostimulation did not lead to the production of heat shock proteins or other heat-related cell distress (*34*).

The dominance in L*_o_* phases observed nevertheless resembles that observed in giant unilamellar vesicles (GUVs) with compositions within the L*_o_*/L*_d_* fluid coexistence region (*44–46*). In particular, we note that just several minutes of SRBW mechanostimulation induces a transition towards a more gel-like phase that is also accompanied by a change in the L*_o_* coverage, primarily due to fusion or coalescence of the L*_o_* domains (*47, 48*), as seen in Fig. 2b; this is in contrast to the slow coalescence observed for mechanostimulation at 1 MHz or below, wherein over 45 mins of continuous exposure was required, and in which no appreciable morphological changes were observed (*24*). More importantly, we observe from Fig. 2b,d, control over the membrane fluidity with the SRBW power, with powers below 1 W leading to a decrease in the L*_o_* domains and hence a transition in the membrane towards a liquid-like state, whereas higher powers beyond 1 W yielding an increase in the L*_o_* domain coverage and a transition in the membrane towards a gel-like state; similar manipulation of the microdomains can be achieved through the post-exposure incubation duration, as shown in Fig. 2c,e. While mechanical cues such as shear, compression and tension have been reported to facilitate shifts in membrane packing and order, such rapid, direct control over the microdomain organisation has only been shown to date in GUVs (*10*), and not in more complex lipid systems such as cells.

In any case, given the interconnectivity of the plasma membrane with the cytoplasm through the membrane skeleton (*49*), the SRBW-driven alteration of the lipid microdomains is then anticipated to drive reorganisation of the underlying cytoskeletal network. In particular, alterations in the L*_o_* domains or cholesterol content in the plasma membrane manifest in cytoskeletal docking, and thus changes to cell stiffness and morphology. Such changes have previously been observed with SRBW mechanostimulation of both MSCs and endothelial cells, wherein rearrangement of the cytoskeletal structure, namely actin stress fibre formation and remodelling (*17, 19*). In particular, the mechanoresponse to the SRBW forcing follows the aforementioned challenge and recovery phases: upon stimulation (i.e., the initial sonochallenge phase), cells were observed to become smaller, stiffer and rounder, followed by a latent response (within 20–60 mins; i.e., the subsequent sonotransformed phase) in which the cells relax to their ground state. The role of actin in facilitating membrane lipid compartmentalisation is prominent in both cases. In the former sonochallenge phase, the increase in the actin-associated L*_o_* domains not only leads to more rigid cells; transient coalescence of these (i.e., the L*_o_*) domains leads to fusion of L*_d_* domains that lack actin association (*50–53*), inducing the membrane to invaginate and to facilitate recruitment of endosomal sorting complexes required for transport (ESCRT) that is responsible for exosome biogenesis (i.e., the formation of intraluminal vesicles (ILVs) and multivesicular bodies (MVBs)) (*54–57*), which we had previously observed with SRBW mechanostimulation (*15*). Further, ESCRT signalling is also responsible for triggering the relaxation of the microdomains towards their ground state in the latter sonotransformation phase. The MVB docking and fusion with the membrane which occurs as a consequence resulting in the observed enhancement in exocytosis (*15*) that, in turn, relieves the membrane tension; the 30–60 min window over which enhanced exosome secretion is reported (*15*) strongly corroborating with increases in the extracellular vesicle structures observed in the SRBW-excited cells (Fig. S1a), as well as the time over which the plasma membrane returns to its ground state.

### Lipid raft association and piezo channel activation

To complete the picture on how cells respond to the SRBW (and SAW) mechanostimulation— which we have shown above to drive deformations in the plasma membrane leading to its decrease in tension, and, in that process, to facilitate an increase in membrane order and polarisation, we now show how such reorganisation in the microdomains can lead to the myriad of downstream cell fates observed (e.g., cell adhesion, migration and proliferation, vesicle trafficking and exosome biogenesis, immune cell activation, stem cell differentiation, and endothelial barrier modulation) in which the central role of second messenger Ca^2+^ signalling has been implicated. Figure 3a,b, in particular, shows the lateral distance along the plasma membrane between the glycosphingolipids (in particular, monosialotetrahexosylgangliosides (GM1) stained with choleratoxin B (CtxB) as a lipid raft marker) and the piezo channels (in particular, Piezo1), which are mechanically activated ion channels known to facilitate Ca^2+^ transport and to hence initiate its signalling cascades. More specifically, we observe the tendency toward co-localisation of gangliosides and Piezo1 following SRBW mechanostimulation as illustrated in Fig. 3e, as evident by the increasing proximity between the lipid rafts (i.e., the L*_o_* microdomains) and Piezo1 channels (Fig. 3b), which were originally distributed throughout the cells, wherein both can be seen to move towards and concentrate within the nuclear region. As observed previously, however, this effect is also transient, with both the lipid rafts and Piezo1 channels returning towards the cell periphery within 60 mins, and hence divaricating back towards the average equilibrium separation distances seen for cells in their initial ground state as well as that for the untreated (control) cells. We note that this recovery time scale corroborates previous observations of the role of another second messenger, cyclic adenosine monophosphate (cAMP), that has also been identified as an instigator of key signalling pathways associated with the SRBW mechanotransduction dynamics, wherein elevated intracellular cAMP levels were recorded within 30 mins post-incubation following SRBW excitation (*19*).

**Figure 3.**
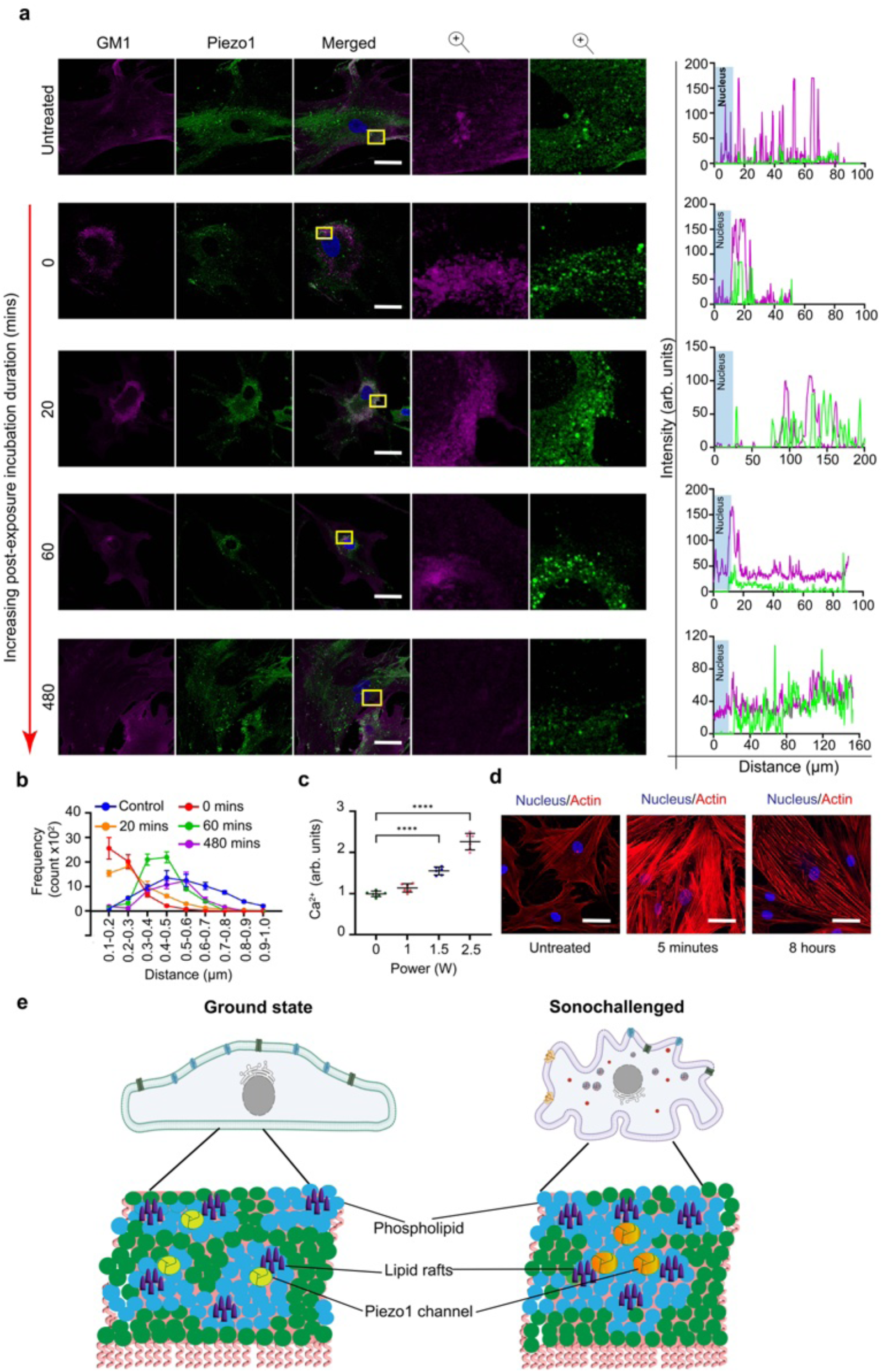
*(preceding page)*: **Lipid raft and Piezo1 reorganisation under the SRBW forcing**. **a** Representative images depicting the distribution of lipid raft residing gangliosides (GM1) and Piezo1 channels in untreated (control) and SRBW-irradiated (2.5 W) cells at different postexposure incubation periods. Cell nuclei were stained using NucBlue™ Live ReadyProbes™ (blue), GM1 with Alexa Fluor 647 conjugate (purple), and, Piezo1 with anti-Piezo1 antibody and Alexa Fluor 488 secondary antibody (green). The third column comprises a merger of the channels with scale bars denoting 50 µm lengths, whereas the last two columns comprise enlarged views of the region indicated in the merged channel. The corresponding line intensities of the GM1 (purple) and Piezo1 (green) fluorescent signals with respect to that for the nucleus (blue) is shown on the right. **b** Frequency distribution (n = 150) of the distance between GM1 and Piezo1 in untreated and SRBW-irradiated (2.5 W) cells at different post exposure incubation times. **c** Intracellular Ca^2+^ (n = 6) as a function of the SRBW power. **d** Images showing actin reorganisation in untreated and SRBW-irradiated cells (2.5 W for 5 mins) at different post-exposure incubation times; the actin structures were stained with ActinRed™ 555 ReadyProbes™ (red) and cell nuclei were stained with NucBlue™ Live ReadyProbes™ (blue); the scale bars denote a length of 50 µm. **e** Schematic illustration showing redistribution of the lipid raft and Piezo1 channels under the SRBW forcing: untreated cells or those initially at ground state prior to the SRBW irradiation comprised a heterogeneous microdomain composition, whereas sonochallenged cells immediately following SRBW irradiation displayed increasing homogeneity in L*_o_* microdomains, whose coalescence leads to their clustering with membrane-associated proteins such as Piezo1, as evident by their increased co-localisation, which, in turn, results in Piezo1 activation and an increase in intracellular Ca^2+^ into the cell. The data are represented in terms of its mean *±* the standard error over triplicate runs, and the asterisks *^⇤⇤⇤⇤^* indicate statistically significant differences with p < 0.0001, respectively.

Additionally, our observations potentially shed light on a possible reason MscL, Piezo1 and Transient Receptor Potential A1 (TRPA1) channels—membrane-associated mechanosensitive ion channels similar to Piezo1—have been reported to be activated at 30 and 7 MHz (*29,58,59*), respectively, the latter being shown to confer sensitivity of neuronal cells to mechanostimulation at that frequency.

As illustrated in Fig. 4a, we thus postulate the chain of events triggered by the SRBW mechanostimulation that leads to activation of the Piezo channels, and subsequently Ca^2+^ mobilisation within the cell to result in the observed biological fates, as follows. Dynamic changes in membrane tension as a result of the deformation to the plasma membrane induced by the SRBW forcing leads to transient increases in membrane order and polarisation (i.e., increasing L*_o_*/L*_d_* ratio), as manifested by the coalescence of the lipid rafts toward the nuclear regions and away from the L*_d_* regions along the cell periphery (Fig. 3a). This provides an environment for their clustering with membrane-associated proteins, such as glycophosphatidylinositol (GPI)anchored proteins and mechanosensitive ion channels such as Piezo1 (Fig. 3e). Activation of these piezo channels is then a direct consequence of this crowding effect—only hypothesized theoretically to date (*60*), together with changes in the membrane curvature and tension directly due to the SRBW-driven membrane deformation, through a Force-from-Lipid model (*61, 62*). Simultaneously, anchoring of cytoskeletal actins to the lipid rafts via the membrane skeleton facilitates concurrent remodelling of the actin cytoskeletal network during SRBW microdomain reorganisation (Fig. 3d), which also allows for activation of the Piezo channels through a Force-from-Filament model (*61, 62*). Together, the concomitant modulation of Ca^2+^ into the cell (Fig. 3c) then allows for triggering of various downstream signalling cascades, such as the ALIX-mediated ESCRT, Rho–ROCK, and cAMP-mediated Epac–Rap1 pathways to result in the various downstream fates observed (*11, 15, 17, 19*).

**Figure 4:**
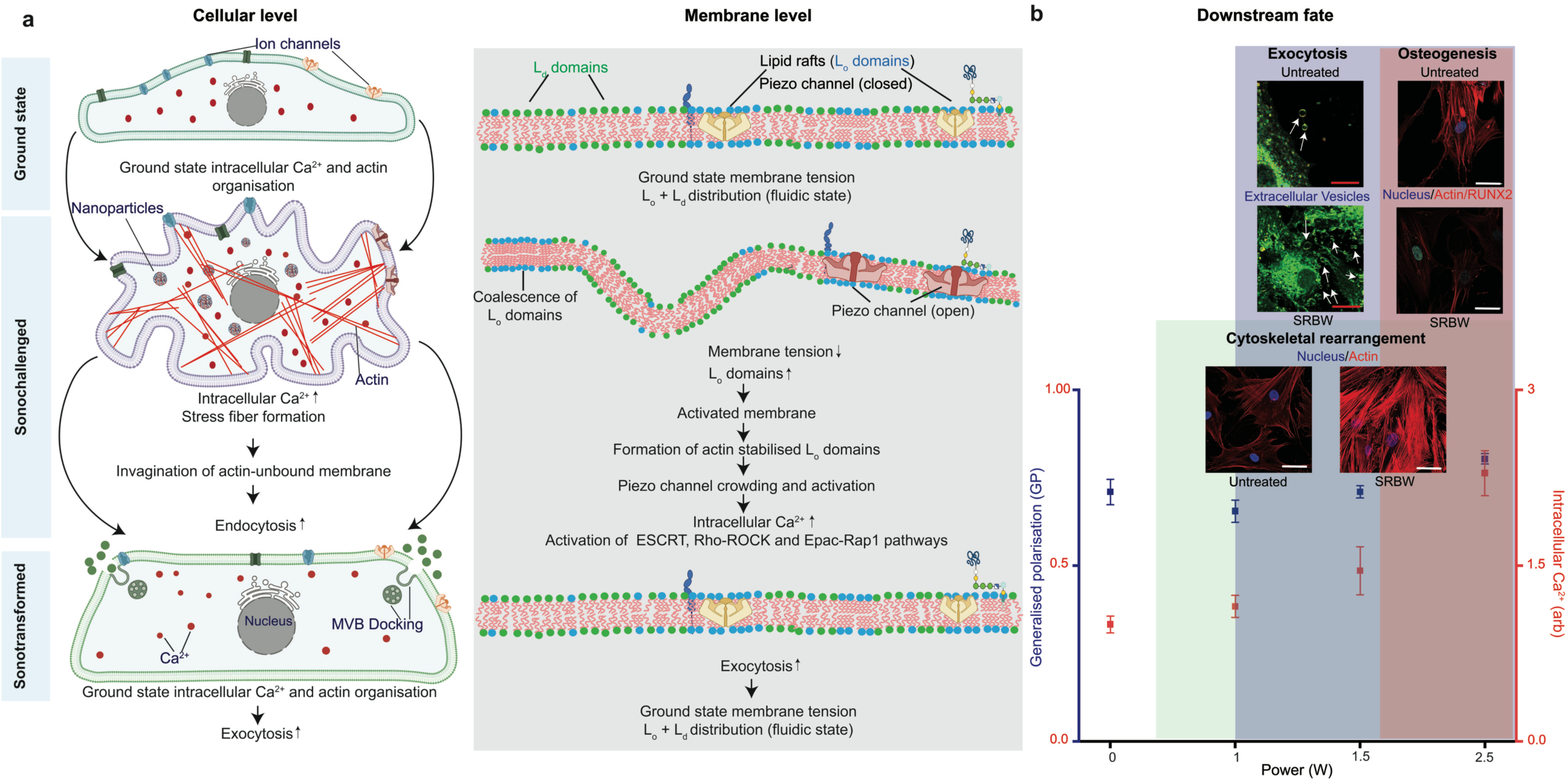
High frequency mechanotransduction and downstream cell fate. a Schematic illustration of the fundamental cellular (left) and (membrane) level mechanisms that govern the SRBW mechanotransduction process to yield diverse downstream cell fates that are b dependent on the modulation of Ca2+ into the cell with the SRBW power, which, in turn, is strongly correlated with the Lo/Ld ratio, as represented by the GP value associated with the membrane lipid order probe C-Laurdan with which the hMSCs are stained. A slight shift in lipid order at input powers between 0.5 and 1.5 W can be seen to initiate cytoskeletal reorganisation, which, together with a 1.5-fold enhancement in intracellular Ca2+ triggers appreciable increase in the biogenesis and exocytosis of extracellular vesicles (indicated by the arrows) due to initiation of the ESCRT pathway. At higher powers, the increase in Lo microdomains, and their subsequent coalescence, yields a 2-fold increase in intracellular Ca2+ that directs the hMSCs along an osteogenic differentiation pathway. The scale bars denote lengths of 20 µm (exocytosis image) and 50 µm (cytoskeletal rearrangement and osteogenesis images).

### Manipulation of lipid microdomains enables control of membrane–protein interactions and downstream cell fate

Figure 4b shows that the downstream cell fate as a consequence of the SRBW mechanostimulation is connected to its ability to manipulate and hence spatiotemporally control lipid microdomain organisation. More specifically, the different cell fates, namely exocytosis (exosome biogenesis and secretion), actin cytoskeletal remodelling and osteogenic differentiation can be seen to be directed simply by varying the input SRBW power, given its ability to modulate the L*_o_*/L*_d_* ratio (as represented by the GP values) and hence the intracellular Ca^2+^ concentration—a consequence of the activation of Piezo1, as discussed in the preceding section—that orchestrates the triggering of various different signalling cascades, such as the ESCRT and Rho–ROCK-and Epac–Rap1 pathways.

To further highlight the potential implications of such direct spatiotemporal control over the lipid microdomains, we also demonstrate here the transformation of tau—a microtubule-binding intrinsically disordered protein, whose significance is pronounced under pathological conditions associated with neurodegenerative diseases—under the high frequency SRBW mechanostimulation. Figure 5, in particular, shows the concentration of tau proteins along with lipid raft fusion (as evident from the higher L*_o_*/L*_d_* ratio in Fig. 5a–c, similar to that observed with the hMSCs in Fig. 2) in SH-SY5Y neuroblastoma cells near the cell nucleus when exposed to the SRBW mechanostimulation (Fig. 5d); these changes being dependent on the membrane composition, cholesterol content and cytoskeletal structure induced by the SRBW forcing, all of which influence the modulation of Ca^2+^ levels in the cell. Consistent with our previous observations, we also note the reversibility of the process, given the tendency for the tau protein to reorganise and the membrane to revert to its ground state after 15–30 mins following cessation of the mechanostimulation. These results are further confirmed by the inhibitor studies in Fig. 5e, where suppression of the SRBW-induced changes in the tau dynamics was observed in the presence of methyl-p’-cyclodextrin to deplete the cholesterol content in the cell, and the membrane permeable intracellular Ca^2+^ chelator bis-acrylamide, 1,2-bis(2-aminophenoxy)ethane-N,N,N’,N’-tetraacetic acid (BAPTA-AM), which acts to deplete the intracellular calcium store. We note that such accumulation of tau in the L*_o_* domains (the reduction in tau expression being attributed to SRBW-induced exocytosis), and their binding to GM1 and phosphatidylserine in the lipid rafts (Fig. 5d), along with accompanying changes in cholesterol, intracellular Ca^2+^ and the corresponding cytoskeletal remodelling, has been pathologically observed during many cases of neurodegeneration (*63–66*), where it has been shown that these membrane–tau interactions are accompanied by increased toxicity (*27, 28, 67*). In any case, this demonstration of membrane–tau interactions under the high frequency mechanostimulation then attests to the amenability of the SRBW platform as a facile method for studying the membrane association dynamics of intrinsically disordered proteins, and, in doing so, opening up the possibility for understanding the complex cellular processes that drive tau dynamics under normal and pathological conditions, which, thus far, has been limited given that such studies have only been able to be conducted till now on *model lipid bilayers* (i.e., vesicles) (*67–70*).

**Figure 5.**
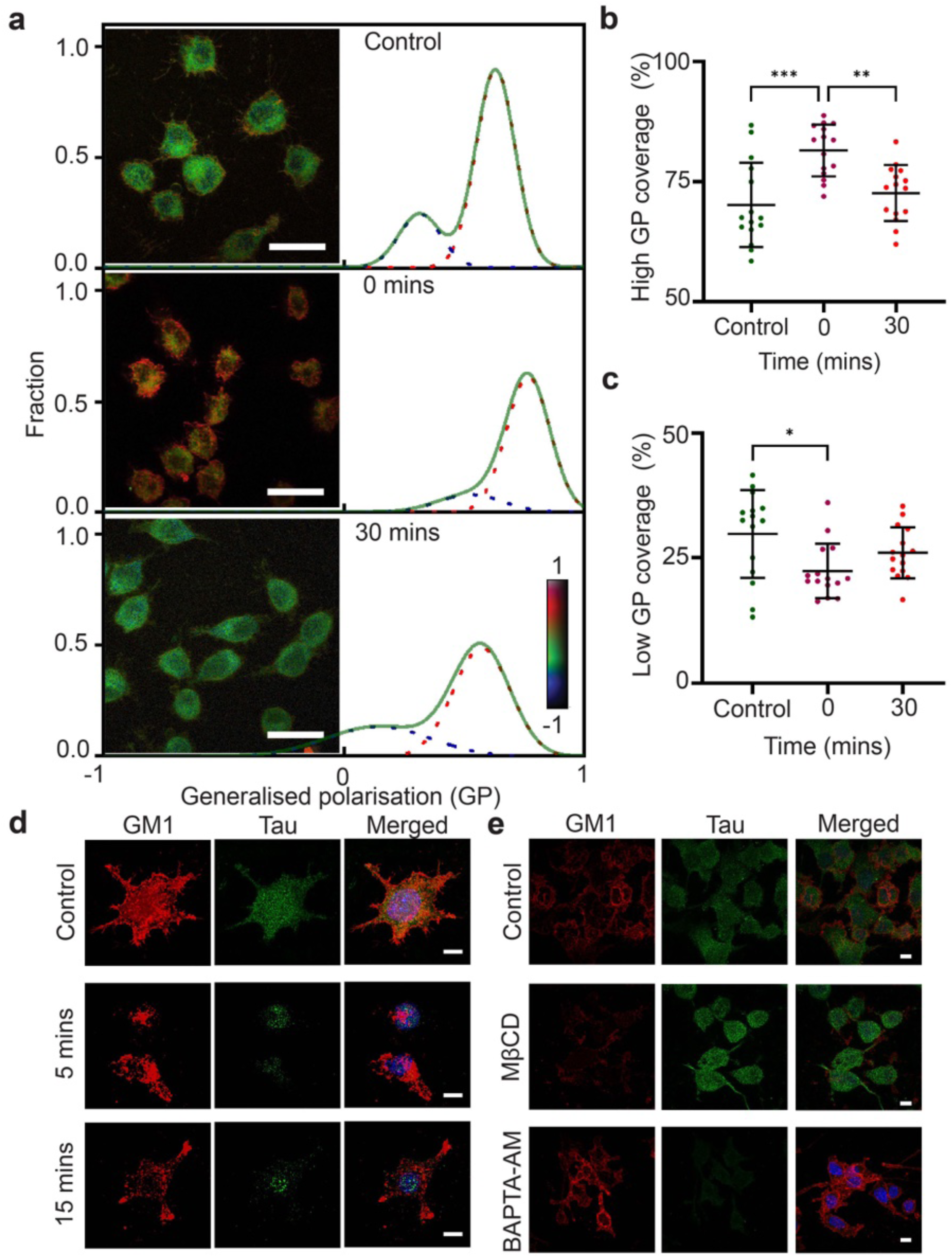
*(preceding page)*: **SRBW induced membrane lipid and membrane-associated protein distribution**. **a** Images (inset) and distribution of their GP values (*-*1 to +1; as represented by the colour bar) for untreated (control) and SRBW-irradiated (2.5 W for 5 mins) C-Laurdan-labelled SH-SY5Y neuroblastoma cells fixed at 0 and mins post-exposure incubation. The histograms for the GP values were fitted with Gaussian distributions, with the solid green curves representing the average between low (dashed blue curves) and high (dashed red curves) distributions. Scale bars denote a length of 30 µm. **b** High and **c** low GP coverage of both untreated and SRBW irradiated cells, obtained by resolving the GP histograms in both cases with the two best Gaussian fits. **d** Lipid raft and tau protein organisation in response to SRBW (2.5 W for 5 mins) at different post-exposure incubation times, and, **e** in the presence of BAPTA-AM and methyl-p’-cyclodextrin (Mp’CD); the control comprised cells unexposed to SRBW stimulation but incubated with media containing the same concentration in DMSO. Images were acquired at 100*⇥* and 60*⇥* magnification, respectively, and the scale bars denote 10 µm lengths. Cell nuclei were stained using NucBlue™ Live ReadyProbes™ (blue), tau proteins with anti-tau antibody and Alexa Fluor 488 secondary antibody (green), and, GM1 with Alexa Fluor™ 647 conjugate (red). The column on the right comprises a merger of these two channels. The data are represented in terms of its mean *±* the standard error over triplicate runs, and the asterisks *^⇤^*,*^⇤⇤^* and *^⇤⇤⇤⇤^* indicate statistically significant differences with p < 0.05, p < 0.01 and p < 0.0001, respectively.

## Conclusion

By correlating changes to the membrane tension, and consequently, the plasma membrane lipid structure, to deformations to the cell imposed by high frequency (10 MHz) mechanostimulation in the form of SRBWs, we have not only been able to explicate the fundamental mechanisms that explain the peculiar and unexpected response of cells to high frequency MHz-order mechanostimulation, but to also show, in addition, the possibility for direct controlled manipulation of lipid microdomains—without necessitating biochemical factors.

In particular, we show that decreases in the deformation-induced membrane tension as a consequence of the SRBW excitation lead to remodelling of the membrane lipid microdomain due to alterations in the membrane fluidity: with increasing SRBW power, we observe an increase in microdomain ordering (i.e., increase in L*_o_*/L*_d_*) and packing, which is accompanied by coalescence of the ordered L*_o_* microdomains and hence a decrease in their size. This has several consequences: (1) due to their stiffer gel-like state compared to the fluid-like L*_d_* microdomains, there is a concomitant shrinkage in the cell dimensions; (2) given the interconnectivity of the membrane skeleton with the cytoskeleton, the underlying actin cytoskeletal network is also altered; (3) clustering of lipid rafts rich in transmembrane proteins, leading to increased proximity and co-localisation of ordered membrane-associated proteins (such as the mechanosensitive ion channels) with the L*_o_* microdomains, results in a crowding effect that is responsible for piezo channel activation (which has only been predicted theoretically to date (*60*)), which, in turn, mobilises influx of the second messenger Ca^2+^ into the cell to trigger a slew of signalling cascades (e.g., ESCRT, Rho–ROCK and Epac1–Rap1) to direct the various downstream cell fates observed. These observations, together with the direct correlation between the L*_o_*/L*_d_* ratio with the SRBW input power, therefore explicitly implicate the role of microdomain organisation (and, more broadly, membrane order and polarisation) on the determination of downstream cell fate.

In addition to the ability of the SRBW-driven spatiotemporal microdomain manipulation to influence intrinsically *ordered* membrane-associated proteins (e.g., Piezo1), we also show its potential for transformation of intrisically *disordered* membrane-associated proteins, namely, tau proteins. Given that the aggregation of tau proteins and their association with the lipid membrane has been implicated in various neurodegenerative diseases, this first demonstration of such a unique possibility highlights the potential of the SRBW mechanostimulation platform as a facile tool to study membrane–tau interactions and the complex signalling cascades resulting from their association. We therefore envisage the platform to facilitate more thorough investigations into the transformation of normal cells towards pathological conditions, and, in doing so, aiding the development of therapeutics to treat neurodegenerative diseases—an endeavour that has currently been limited due to the lack of a means for directly manipulating membrane microdomain organisation in systems such as cells that are considerably more complex than the model bilayer systems that such manipulation has typically been demonstrated on to date.

## Materials and Methods

### Materials

Human mesenchymal stem cells (hMSCs) were acquired from Lonza Pty. Ltd. (Mount Waverley, VIC, Australia), whereas the SH-SY5Y neuroblastoma cell line was obtained through Dr Shwathy Ramesan at The Florey Institute of Neuroscience and Mental Health (Parkville, VIC, Australia). Silicone oil, Triton™ X-100, methyl-p’-cyclodextrin (Mp’CD), dimethylsulphox- ide (DMSO), Opti-MEM™ phosphate buffered saline (PBS), Gibco penicillin–streptomycin, trypsin–ethylenediaminetetraacetic acid (EDTA), formaldehyde, bovine serum albumin (BSA), fetal bovine serum (FBS), Dulbecco’s Modified Eagle Medium (DMEM), Eagle’s Minimum Essential Medium, F12 Medium, Fura-2 acetoxymethyl ester (Fura-2AM),1,2-bis(2-aminophenoxy) ethane-N,N,N’,N’-tetraacetic acid tetrakis(acetoxymethyl ester) (BAPTA-AM), Trypan Blue solution, cholera toxin subunit B (recombinant; CtxB), ActinRed™ 555 ReadyProbes™, NucBlue™ Live ReadyProbes™, Alexa Fluor 647 conjugate, anti-Piezo1 antibody, T25 cell culture flasks and Nunc™ Lab-Tek™ II Chambered Coverglass were sourced from Thermo Fischer Scientific Pty. Ltd. (Scoresby, VIC, Australia). FluoroDish cell culture plates were procured from World Precision Instruments (WPI) Ltd. (Hertfordshire, England). Anti-tau rabbit monoclonal antibody, and, anti-rabbit and anti-mouse IgG (H+L) F(ab’)2 Fragment (Alexa Fluor 488 conjugate), on the other hand, were obtained from Cell Signaling Technology Inc. (Danvers, MA, USA), whereas C-Laurdan and Flipper-TR™(CYT-CY-SC020) were obtained from R&D Systems (Minneapolis, MN, USA) and Cytoskeleton Inc. (Denver, CO, USA), respectively.

### Device fabrication

The SRBW devices shown in Fig. 1a consisted of 40 alternating pairs of 11-mm-wide and 66-nm-thick straight interdigitated aluminium transducer (IDT) electrodes arranged in a basic full-width interleaved configuration on 500-µm-thick 127.86*^o^* Y –X rotated lithium niobate (LiNbO_3_) single-crystal piezoelectric substrates (Roditi Ltd., London, UK). Sputter deposition and standard ultraviolet (UV) photolithography were used to pattern the IDTs on top of a 33- nm-thick chromium adhesion layer. The resonant frequency of the device f = c/A where c is the SRBW phase speed in LiNbO_3_, was set at 10 MHz by prescribing the width and gap of the IDT fingers (A/4). To generate the SRBW, a signal generator (SML01; Rhode & Schwarz Pty. Ltd., North Ryde, NSW, Australia) and amplifier (10W1000C; Amplifier Research, Souderton, PA, USA) were used to apply an alternating electrical signal to the IDT at the resonant frequency. The acoustic wave energy from the device was transmitted to the cells adherent to the glass-bottom chamber slide through a thin layer of silicone oil with a viscosity of 45–55 cP and density 0.963 g/ml at 25 *^o^*C.

### Cell culture and mechanostimulation

The hMSCs were grown in DMEM with 1% penicillin–streptomycin and 10% FBS in a humid- ified incubator at 37 *^o^*C and 5% CO_2_ until they covered 80-90% of a standard T25 flask. They were then detached using 0.05% trypsin–EDTA and transferred to 8-well plates or FluoroDish cell culture plates at a seeding density of 3,000 cells per well before being incubated for 28 hrs in DMEM to ensure proper adhesion. SH-SY5Y neuroblastoma cells, on the other hand, were expanded in a 1:1 mixture of Eagle’s Minimum Essential Medium and F12 Medium with 10% FBS and 1% penicillin–streptomycin.

The cells in the well-plate were subsequently exposed to 10 MHz SRBW irradiation at different input powers and exposure times. Unless otherwise specified, the exposure time was 10 mins for input powers below 2 W, and 5 mins for input powers of 2 and 2.5 W. The cells were incubated immediately for different periods following cessation of the SRBW irradiation before being processed for further analysis. Control samples consisted of cells seeded in DMEM with 1% penicillin–streptomycin and 10% FBS at the same density and incubated for the same time period, but without any vibrational excitation.

For the inhibitor studies, the SHSY-5Y were seeded at 3,000 cells per well and incubated with BAPTA-AM (10µM, for 25 mins) or Mp’CD (300 µM, for 60 mins) prior to SRBW irradiation and washed with media devoid of the inhibitors. The irradiated cells were then fixed after 5 mins of post-exposure incubation. The control comprised cells unexposed to SRBW stimulation but incubated with media containing the same concentration in DMSO.

### Membrane tension

The process for characterising the membrane tension of the cells is as follows. The cells were first incubated with 3 µM Flipper-TR™ in DMSO for 15 mins at 37 *^o^*C, and subse- quently washed thrice in media devoid of the stain. The cells were then exposed to the SRBW mechanostimulation at different input powers (1.5 and 2.5 W), following which they were fixed at various post-exposure incubation times. Fluorescence lifetime imaging (FLIM) was subsequently carried out with an inverted confocal laser scanning microscope (FV3000; Olympus Corp., Tokyo, Japan) equipped with a time-correlated single-photon counting module (Pico-Quant GmbH, Berlin, Germany) to measure the lifetime of the probe. The FLIM excitation source comprised a pulsed 485 nm laser (LDH-D-C-485; PicoQuant GmbH, Berlin, Germany) operating at 20 MHz, and the emission signal was collected through a 600/50 nm bandpass filter using a gated hybrid photomultiplier detector (PMA Hybrid 40; PicoQuant GmbH, Berlin, Germany) and a time-correlated single photon counting and multi-channel scaling board (TimeHarp 260 PICO; PicoQuant GmbH, Berlin, Germany). The lifetimes of the probe were then determined from the acquired FLIM images using the supplied software (SymPhoTime 64; Pico- Quant GmbH, Berlin, Germany). The area of the cells and the protrusion-like structures, on the other hand, were measured using ImageJ (National Institutes of Health, Bethesda, MD, USA), the latter using the plugin FiloQuant; examples of how the cell boundaries and protrusions were defined for the analysis using the plugin are shown in Fig. S2.

### Membrane polarisation

Membrane polarisation was assessed by first treating the cells with 3 µM C-Laurdan in DMSO, and washing them three times in media devoid of the stain. The cells were then exposed to the SRBW mechanostimulation at various input powers (0.5, 1, 1.5, 2 and 2.5 W). All cells, except those exposed to 2.5 W, were fixed at the end of the 10-min exposure duration. The cells that were exposed to SRBW at 2.5 W (5 mins) were fixed at various post-exposure periods (0, 20, and 60 mins). Two-photon fluorescence images were captured using a confocal microscope (FV3000; Olympus Corp., Tokyo, Japan) with 780 nm excitation at 2.5% laser power, and using a L100 oil-immersion objective (numerical aperture, NA = 1.30). The intensity images were recorded across two channels with emission in the range of 400–460 and 470–530 nm. The generalised polarisation (GP) values were determined using the ImageJ macro (National Institutes of Health, Bethesda, MD, USA) specified in Ref. (*71*), along with a sensitivity correction factor of 0.3, which was established by measuring the relative sensitivities of the two channels. GP distributions were derived from normalised frequency histograms and fitted to one or two Gaussian functions with a nonlinear fitting algorithm (Origin 7.0; OriginLab Corp., Northampton, MA, USA). Only fits for which the Chi-square test yielded values of p > 0.95 were considered acceptable. The area under the peaks and the maximum peak fit values were respectively used to define the coverage (relative percentage) of the L*_o_* and L*_d_* microdomains, and the high and low GP values for each individual experiment, which was independently replicated at least thrice. Images displayed in the figures are representative of the wider data set that was used for quantification.

### Immunofluorescence staining

The cells requiring fixation were washed thrice in PBS and incubated in 4% formaldehyde for 20 mins at room temperature, followed by three further wash cycles in PBS. The cells were then blocked by incubating them in 5% BSA in PBS for 60 mins, followed by further incubation with CtxB Alexa Fluor 647 conjugate overnight at 4 *^o^*C. The cells were subsequently washed thrice in PBS, permeabilised with 0.1% Triton™ X-100 for 5 mins, washed a further three times in PBS, and then blocked again with 5% BSA in PBS for 60 mins, following which they were incubated with anti-Piezo1 (1:500) or anti-tau (1:500) antibodies overnight at 4 *^o^*C. After three washes in PBS, the cells were next incubated in the secondary antibody (anti-rabbit and anti-mouse IgG (H+L) F(ab’)2 Fragment (Alexa Fluor 488 conjugate); 1:1000) for 1 hr in the dark at room temperature and subsequently washed three more times in PBS. Nuclei were counterstained using NucBlue™ Live ReadyProbes™ while actin filaments were stained with ActinRed™ 555 ReadyProbes™ prior to imaging under a confocal microscope (A1 HD25; Nikon Instruments Inc., Melville, NY, USA). Lipid raft (ganglioside, GM1) and Piezo1 co-localisation distances were measured using the ImageJ distance analysis plugin DiAna (National Institutes of Health, Bethesda, MD, USA), whereas the distribution studies in Fig. 3a were analysed using the plot profile in ImageJ (National Institutes of Health, Bethesda, MD, USA). Figure S2 shows examples of how the cellular boundaries and protrusion-like features were defined for the analysis.

### Intracellular Ca**^2+^**

The fluorescent calcium indicator Fura-2AM was used to measure free cytosolic Ca^2+^ levels. Briefly, the cells were placed in 5 µmol/l Fura-2AM in Opti-MEM™ reduced serum medium with 2% (vol/vol) heat-inactivated FBS and incubated at 37 *^o^*C in the absence of light. After 1 hr of incubation, the medium with extracellular Fura 2AM was replaced with fresh medium followed by further incubation for an additional 20 mins before irradiating the sample with the SRBW. Using a spectrophotometric plate reader (CLARIOstar, BMG LabTech, Mornington, VIC, Australia), we subsequently measured the fluorescence emission intensity at 510 nm in individual wells at excitation wavelengths of 340 and 380 nm. The ratio of Fura-2AM fluorescence emission in response to 340 nm and 380 nm excitation (340/380) can then be calculated and expressed as the fold change relative to that measured for the respective control (unexcited) cells.

### Statistics

Quantitative data were presented as the mean *±* standard deviation from a minimum of three independent experiments. Statistical analysis involved one-way analysis of variance (ANOVA); a threshold of p < 0.05 signified statistical significance.

## Acknowledgments

LYY and CEC acknowledge support from the Australian Research Council through Discovery Project grant DP210101720.

## Supplementary Materials

Figs. S1 and S2

## Footnotes

* Most studies, in fact, have attempted to demonstrate that ultrasound does not cause any noticeable effects on cells (12).

† Lower GP values are typically associated with the dominance of Ld domains wherein the plasma membrane has a closer resemblance to a fluid-like state, whereas higher values are typically associated with the dominance of Lo domains in which the plasma membrane resembles a stiffer gel-like state (40–43) (Fig. 2a).

