## Supplementary Figures for "High Frequency MHz-Order Nanovibration Enables Cell Membrane Remodelling and Lipid Microdomain Manipulation"

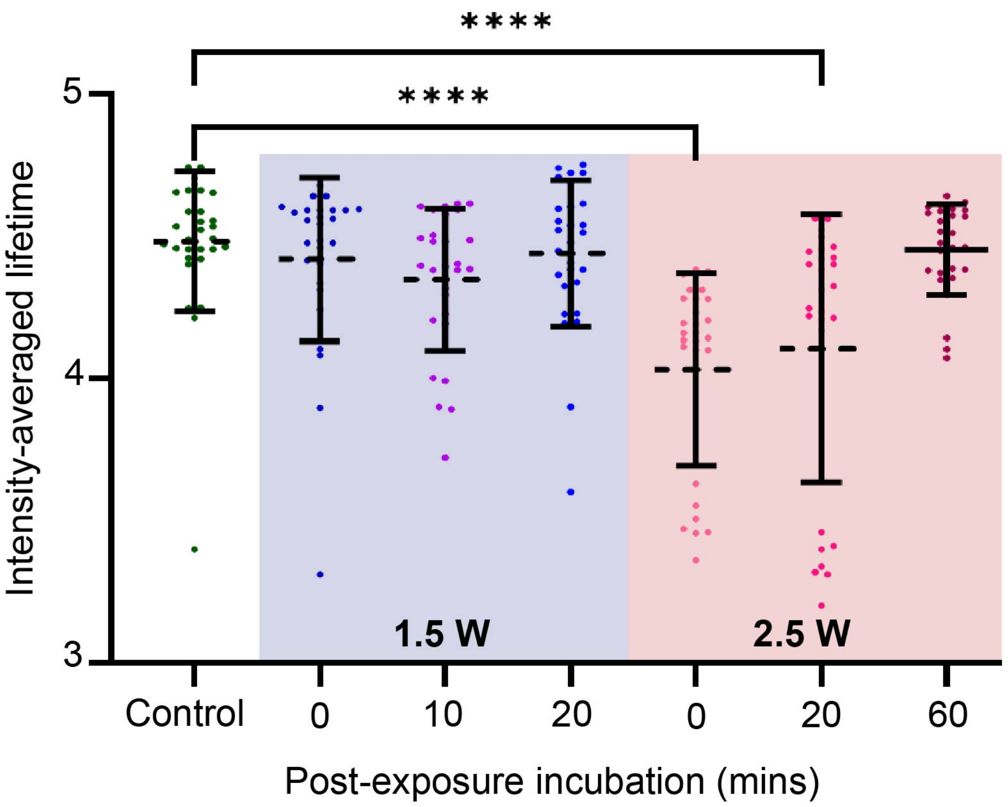


**Figure S1** Intensity-averaged fluorescence lifetime as a function of the SRBW input power and post -exposure incubation duration. The data are represented in terms of its mean ± the standard error over triplicate runs, and the asterisks **** indicate statistically significant differences with $p < 0.0001$.

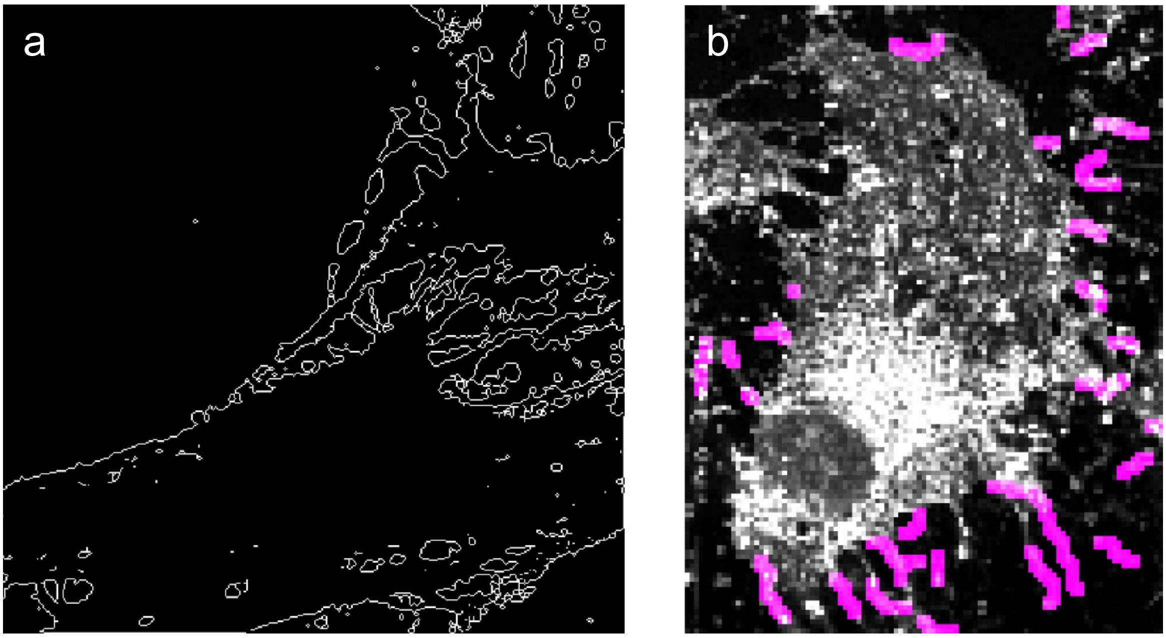


**Figure S2** (a) Example showing how the area of cell is measured using ImageJ in which the outline of the cell is used to define its boundary. (b) Example showing protrusion-like structures identified using FiloQuant plugin in ImageJ. The purple lines were identified as the protrusions and are used to determine their number and length.

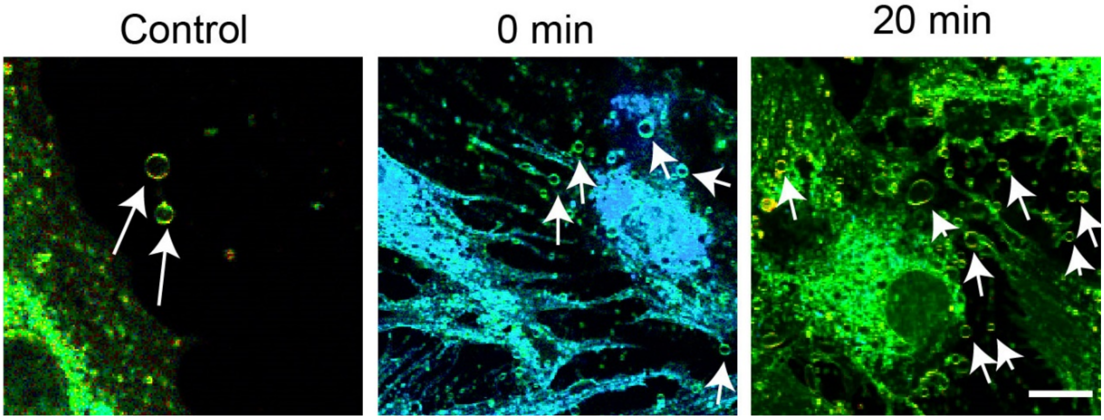


**Figure S3** Microscopy images showing the release of extracellular vesicles, indicated by the arrows, from untreated (control) and SRBW-treated (2.5 W) cells (stained with Flipper-TR$^{TM}$) fixed at different post-exposure incubation (0 and 20 mins). The scale bar denotes a length of 20 μm.
